# Dual targeting of CCR2^+^ monocytes and neutrophils enhances anti-tumor immunity

**DOI:** 10.64898/2026.05.13.723724

**Authors:** Klaire Yixin Fjæstad, Astrid Zedlitz Johansen, Hannes Linder, Marco Carretta, Majken Siersbæk, Kevin James Baker, Marie-Louise Thorseth, Mie Linder Hübbe, Mads Hald Andersen, Lars Grøntved, Daniel Hargbøl Madsen

## Abstract

Targeting immunosuppressive tumor-associated myeloid populations has emerged as a promising strategy to enhance anti-tumor immunity. The CCL2–CCR2 axis plays a central role in the recruitment of monocytes that differentiate into tumor-associated macrophages (TAMs), yet the therapeutic potential of CCR2 targeting remains limited. Using transgenic CCR2-DTR mice, we show that depletion of CCR2^+^ monocytes and TAMs reduced tumor growth across multiple models, accompanied by remodeling of the tumor microenvironment (TME). Residual CCR2-independent TAMs exhibited a pro-inflammatory and less immunosuppressive phenotype, and expressed the alternative recruitment receptor CCR3. Concomitantly, CCR2 depletion markedly enhanced anti-tumor immunity by increasing infiltration of activated CD8^+^ T cells. Splenocytes from tumor-bearing CCR2-DTR mice showed an increased IFNγ response to a cancer-associated antigen. Furthermore, CCR2 depletion synergized with immune checkpoint blockade to enhance tumor control. Despite these effects, compensatory tumor infiltration of neutrophils following CCR2 targeting limited therapeutic benefit. These neutrophils exhibited a terminally differentiated, immunosuppressive phenotype and were associated with increased cancer cell-intrinsic expression of the neutrophil-recruiting chemokines *Cxcl2* and *Cxcl5*. Importantly, combined depletion of CCR2^+^ cells and neutrophils overcame this resistance mechanism, resulting in reduced tumor growth, prolonged survival, and complete tumor clearance in 25% of the mice. Dual depletion of CCR2^+^ cells and neutrophils was also associated with a synergistic increase in circulating CD8^+^ T cells. These findings highlight the dynamic remodeling of the TME upon CCR2 depletion and suggest that combinatorial strategies addressing immunosuppressive neutrophil infiltration may improve the efficacy of CCR2 targeting therapies.

## INTRODUCTION

Despite major advances in cancer therapy, durable clinical responses remain limited for many patients (1,2). Immune checkpoint inhibitors (ICIs) have transformed the treatment landscape across multiple cancer types (3,4). However, a substantial proportion of patients either fail to respond or eventually develop resistance (5), highlighting the need to identify mechanisms of immune evasion and develop novel therapeutic strategies.

The tumor microenvironment (TME) plays a crucial role in shaping therapeutic outcomes (6,7). Myeloid cells, including monocytes and tumor-associated macrophages (TAMs), are key drivers of immunosuppression in the TME (8-10). They can directly inhibit cytotoxic T cell function through contact-dependent mechanisms such as the PDL1-PD1 interaction, or promote tumor progression through the secretion of anti-inflammatory cytokines and the recruitment of other immunosuppressive cell types (10,11). Consequently, the abundance of TAMs is associated with poor prognosis in most cancer types (12,13). High levels of circulating monocytes and serum levels of TAM recruitment chemokines also correlate with disease progression in sarcoma and mesothelioma (14,15). Main monocyte and TAM targeting strategies include inhibiting recruitment to the tumor, reprogramming towards an inflammatory state, and promoting phagocytosis and antigen presentation (16-18).

The CCL2–CCR2 axis is critical in mediating the recruitment of monocytes to sites of inflammation (19-21). Upon recruitment to the TME, CCR2^+^ monocytes can rapidly differentiate into TAMs (22-24). CCL2 is frequently overexpressed by cancer cells, and increased serum levels of CCL2 are associated with advanced disease stage, higher tumor burden, and reduced overall survival (25,26). This makes CCR2 an attractive therapeutic target for limiting the infiltration of suppressive monocyte and TAM populations. Preclinical studies have demonstrated that blockade of CCR2 or CCL2 can reduce monocyte trafficking and TAM abundance, and enhance anti-tumor immunity (27-29). These findings led to the clinical development of CCR2-blocking antibodies and small-molecule inhibitors. However, blocking CCR2 as a monotherapy has not achieved significant clinical benefit in cancer and other inflammatory diseases (26,30,31). Potential explanations include tumor heterogeneity between patients, TME diversity, and compensatory recruitment pathways (32,33). Therefore, a deeper understanding of how tumors adapt to CCR2 blockade remains critical for the rational design of myeloid-targeted therapies. Defining compensatory immune mechanisms and identifying potential combination strategies may help overcome therapeutic resistance.

Here, we define the immunological consequences of CCR2 targeting in tumor-bearing mice and identify compensatory mechanisms that limit its therapeutic efficacy. CCR2 depletion attenuated tumor growth in multiple murine tumor models by reducing immunosuppressive myeloid cells, and enhancing cytotoxic lymphocyte infiltration and function. We identify compensatory neutrophil recruitment upon CCR2 depletion, which constrains treatment responses. We demonstrate that concurrent targeting of CCR2^+^ cells and neutrophils can overcome this resistance, resulting in enhanced tumor clearance. This study highlights the plasticity of the myeloid compartment following CCR2 targeting and supports combinatorial strategies that target both the CCR2-CCL2 axis and neutrophils to enhance anti-tumor immunity.

## RESULTS

### CCR2+ monocytes and macrophages contribute to tumor progression

To determine whether CCR2^+^ cells contribute to tumor progression, we utilized CCR2-DTR mice, in which diphtheria toxin (DT) enables targeted depletion of CCR2-expressing monocytes. Forty-eight hours after low-dose DT injection in CCR2-DTR mice bearing MC38 colon adenocarcinoma tumors, circulating CCR2^+^ cells were reduced by approximately 50% compared to DT-treated wildtype littermates (Fig. 1A). This was accompanied by a marked decrease in total circulating monocytes (Fig. 1B), confirming effective targeting. CCR2 is also reported to be expressed at low levels on T cells (34). Therefore, we assessed whether DT treatment also affected this cell population. Low-dose DT did not alter peripheral T cell abundance in CCR2-DTR mice (Fig. 1C), indicating that this approach preferentially depletes monocytes while sparing key effector lymphocytes.

**Figure 1.**
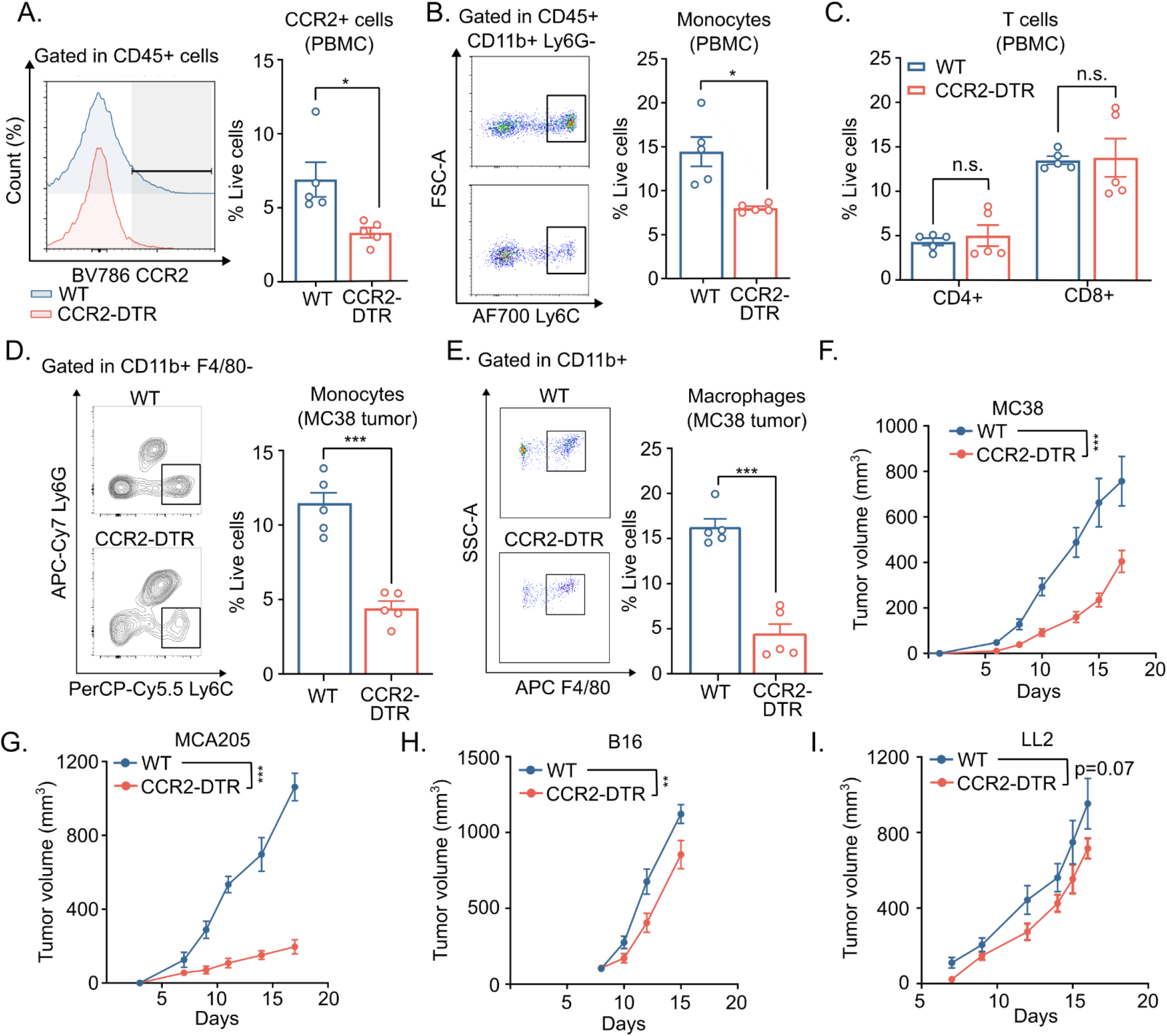
CCR2^+^ cell depletion reduces monocyte and TAM infiltration, and suppresses tumor growth. **(A-C)** Flow cytometry analysis was performed on peripheral blood mononuclear cells (PBMC) in MC38 tumor-bearing mice from wildtype control and CCR2-DTR mice upon diphtheria toxin (DT) treatment. **(A)** Representative flow cytometry plot (left) and quantification of CCR2^+^ cells (right). **(B)** Representative flow cytometry plot (left) and quantification of monocytes (right). **(C)** Quantification of CD4^+^ T cells and CD8^+^ T cells. **(D-E)** Single cell suspensions were made from MC38 tumors at the experimental endpoint and analyzed by flow cytometry. **(D)** Representative flow plot (left), and quantification (right) of monocytes. **(E)** Representative flow plot (left), and quantification (right) of macrophages. Tumor growth of MC38 **(F)**, MCA205 **(G)**, B16 **(H)**, and LL2 **(I)** subcutaneous tumor models in wildtype and CCR2-DTR mice upon DT treatment (n=8-10 per group). Each dot in dot plot represents a biological replicate. n.s. not significant, *p<0.05, **p<0.01, ***p<0.005, according to two-sided multiple t test with Bonferroni correction, or Tum Growth software for tumor growth analyses. Means ± SEM are depicted.

To evaluate how CCR2+ monocyte depletion affected the abundance of monocytes and macrophages in the tumor microenvironment, we inoculated CCR2-DTR and littermate wildtype mice with MC38 cells and treated the mice with DT. Flow cytometry-based analysis demonstrated reduced numbers of tumor-infiltrating monocytes and tumor-associated macrophages (TAMs) in CCR2-DTR mice (Fig. 1D-E). We next evaluated the effect of CCR2^+^ monocyte depletion on tumor progression. CCR2-DTR mice showed significantly reduced MC38 tumor growth (Fig. 1F and Supp. Fig 1A). We tested the effect of CCR2 cell depletion on three other syngeneic mouse tumor models. We observed a similar strong reduction in tumor growth in the MCA205 fibrosarcoma model (Fig. 1G, Supp. Fig. 1B) and a moderate reduction in the B16 melanoma model (Fig. 1H, Supp. Fig. 1C). In the LL2 lung adenocarcinoma model, we observed a trend towards reduced tumor growth upon CCR2^+^ depletion (Fig. 1I, Supp. Fig. 1D).

### CCR3^+^ TAMs sustain an M1-like macrophage population following CCR2 depletion

We next investigated in detail how CCR2 depletion reshapes the tumor myeloid compartment. Although CCR2 depletion reduced the overall abundance of tumor-associated macrophages, a residual macrophage population persisted within the tumors, suggesting the presence of TAMs that are not derived from CCR2^+^ monocytes. We isolated TAMs from MC38 tumors in wildtype and CCR2-DTR mice for whole transcriptome analysis. RNA sequencing of the TAMs showed decreased *Ccr2* expression levels in CCR2-DTR mice, thereby confirming a reduction in CCR2^+^ macrophages in CCR2-DTR mice (Fig. 2A). Overall, transcriptomic analyses revealed markedly distinct gene expression profiles between TAMs from CCR2-DTR mice and wildtype controls (Supp. Fig. 2A-C). To define the phenotype of TAMs, we utilized the transcriptomic deconvolution tool quanTIseq (35). TAMs from CCR2-DTR mice were skewed toward an M1-like phenotype with reduced M2-associated signatures relative to WT controls (Fig. 2B). Consistent with this shift, TAMs from CCR2-DTR tumors showed lower gene expression levels of immunosuppressive molecules, including *Il10* and *Ptgs2* (Fig. 2C).

**Figure 2.**
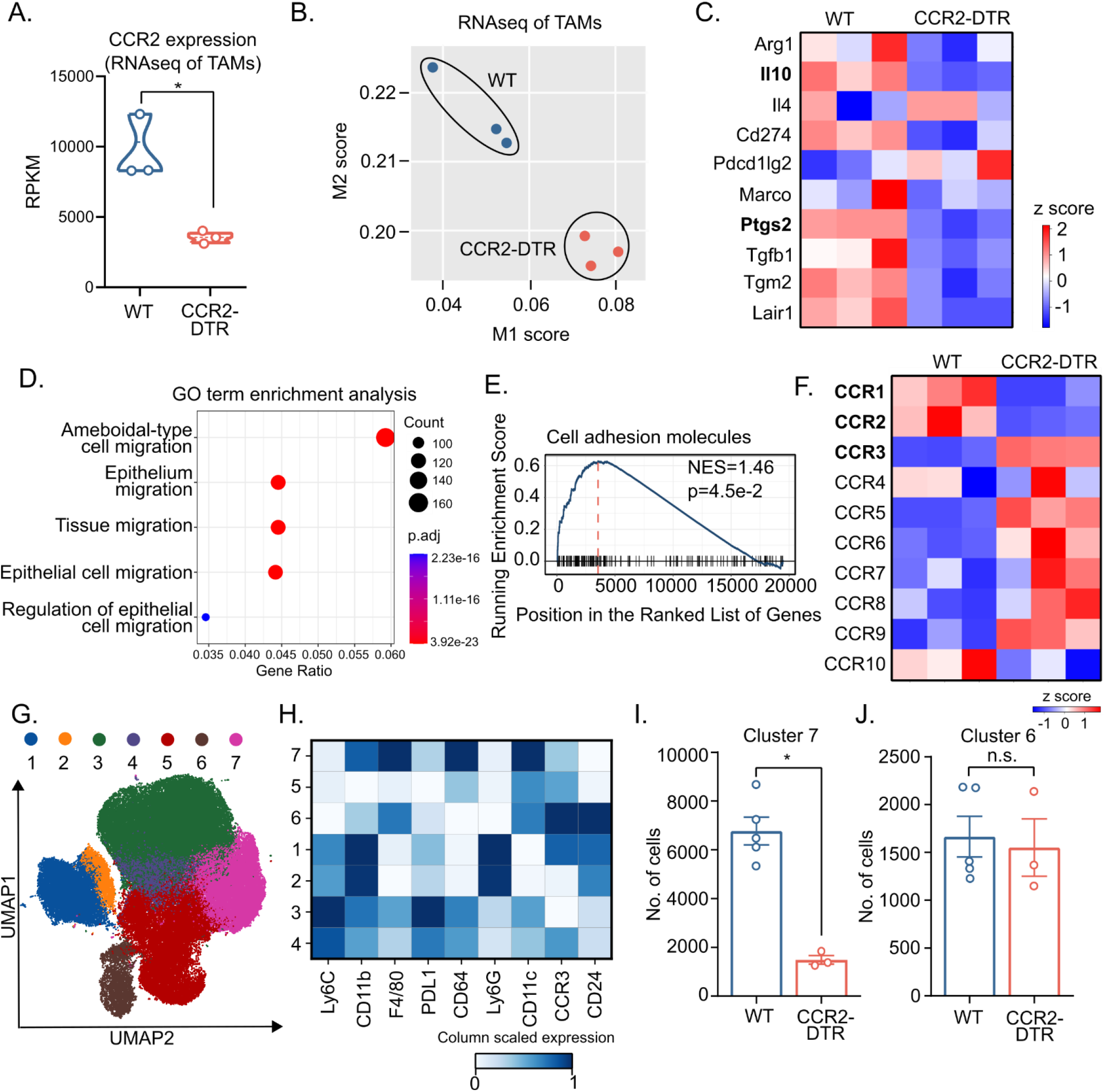
CCR3 mediates CCR2-independent macrophage recruitment. **(A-F)** RNAseq was performed on tumor-associated macrophages (TAM) from MC38 tumor-bearing wildtype and CCR2-DTR mice. **(A)** Expression level of Ccr2 gene. **(B)** QuanTIseq characterization of M1 and M2 macrophage signatures. **(C)** Heatmap of immunosuppressive molecule expression. Bold gene names in heatmaps indicate significantly regulated genes. **(D)** Gene ontology (GO) term enrichment analysis of all upregulated genes in CCR2-DTR mice compared to wildtype controls. **(E)** Plot of Gene Set Enrichment Analysis (GSEA) of KEGG pathways, showing enriched gene set in CCR2-DTR mice. **(F)** Heatmap of CC chemokine receptor expression in TAMs. Bold gene names in heatmaps indicate significantly regulated genes. **(G-J)** Tumor-infiltrating myeloid cells (CD11b^+^ and/or CD11c^+^) from wildtype and CCR2-DTR mice were isolated and stained for high-dimensional flow cytometry clustering (*n*= 3-5 per group). **(G)** UMAP plots showing the distribution of clusters. **(H)** Heatmap of relative expression (column scaled) of markers in myeloid cell FlowSOM clusters. **(I-J)** Absolute cell number in FlowSOM cluster 6 **(I)** and cluster 7 **(J)**. Each dot in dot plot represents a biological replicate. *p<0.05, according to two-sided multiple t test with Bonferroni correction. Means ± SEM are depicted.

Gene ontology (GO) term enrichment analysis of upregulated genes in TAMs from CCR2-DTR mice highlighted pathways related to migration (Fig. 2D), while gene set enrichment analysis of KEGG pathways revealed upregulation of cell adhesion molecule signaling (Fig. 2E). These pathways collectively suggest that residual macrophages in the TME of CCR2-DTR mice may display different migratory behavior and activity. Analysis of a panel of genes encoding CC chemokine receptors also showed distinct expression patterns, and identified CCR3 as significantly upregulated in CCR2-DTR TAMs (Fig. 2F).

To further characterize the myeloid landscape in the TME, we performed high-dimensional flow cytometry profiling of tumor-infiltrating myeloid cells from WT and CCR2-DTR tumors and analyzed the data using FlowSOM clustering. This analysis identified seven distinct clusters, including two monocyte populations (cluster 3 and 4), two macrophage populations (clusters 6 and 7), and two neutrophil populations (clusters 1 and 2) (Fig. 2G-H). Both monocyte populations were profoundly reduced upon CCR2^+^ cell depletion (Supp. Fig. 2E-F). Among macrophages, cluster 7 was selectively depleted, consistent with efficient DT-mediated targeting (Fig. 2I), whereas cluster 6, characterized by high CCR3 expression, was unaffected (Fig. 2J).

### CCR2 depletion enhances lymphocyte infiltration

To examine whether CCR2 depletion modulates key effector cell populations involved in anti-tumor immunity, we analyzed tumor-infiltrating lymphocytes by flow cytometry. CCR2-DTR tumors showed a trend toward increased infiltration of CD8^+^ T cells and CD4^+^ T cells (Fig. 3A-C), without an increase in Tregs (Supp. Fig. 3A). Similarly, we observed an increase in CD4^+^ and CD8^+^ T cells in the MCA205 model upon CCR2 depletion (Supp. Fig. 3B-E). Tumor-infiltrating CD8^+^ T cells displayed an activated phenotype in CCR2-depleted mice, marked by elevated PD1 expression (Fig. 3D). No difference in PD1 was observed in CD4^+^ T cells (Fig. 3E).

**Figure 3.**
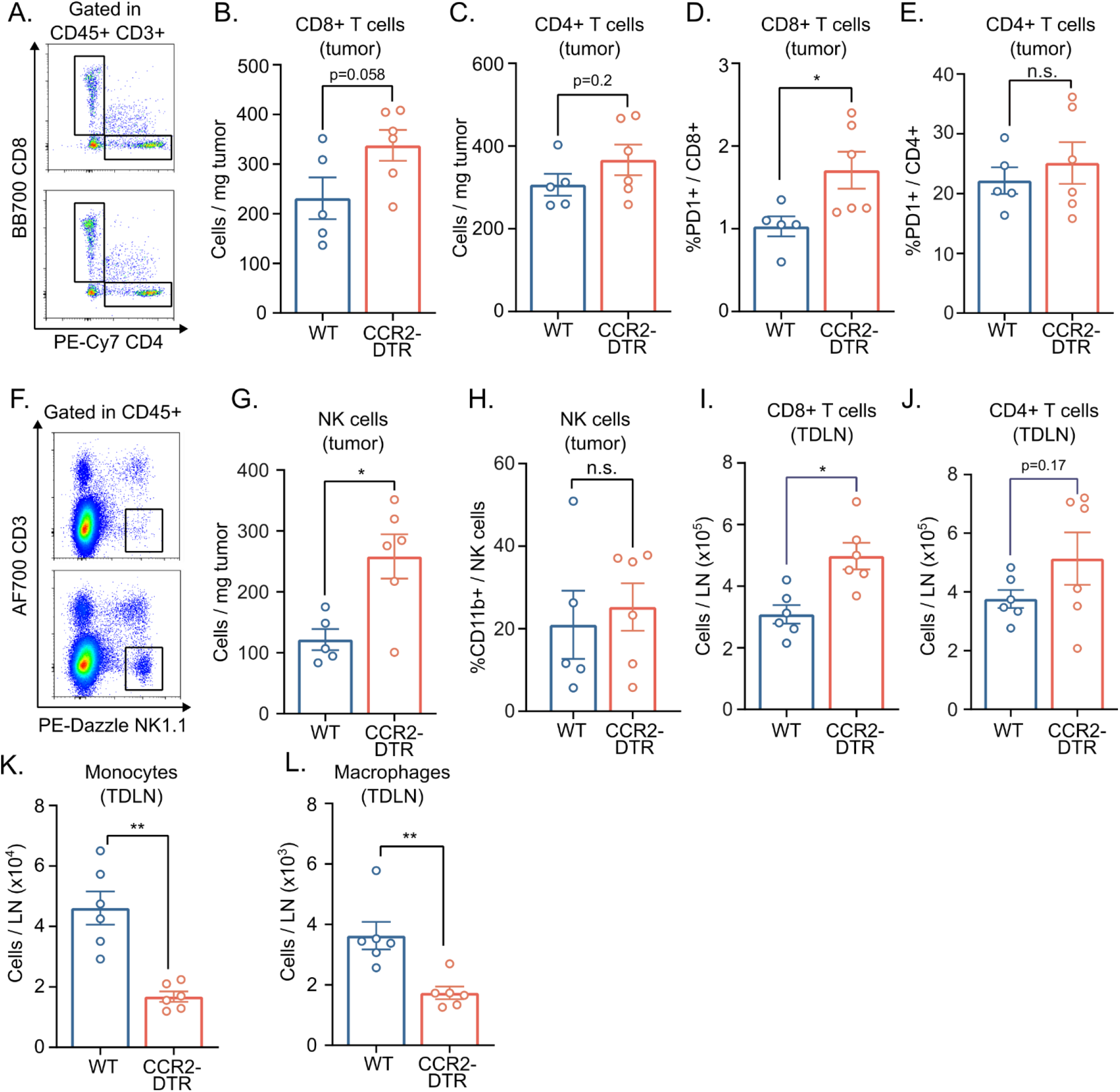
CCR2 depletion enhances lymphocyte infiltration. **(A-H)** Flow cytometry analysis was performed on single cell suspension of MC38 tumors from wildtype and CCR2-DTR mice. Representative flow cytometry plot **(A)** and quantification of absolute cell number of **(B)** tumor-infiltrating CD8^+^ T cells. **(C)** Quantification of absolute cell number of tumor-infiltrating CD4^+^ T cells. PD1 expression on tumor-infiltrating CD8^+^ T cells **(D)** and CD4^+^ T cells **(E)**. Representative flow cytometry plot **(F)** and quantification of absolute cell number of **(G)** tumor-infiltrating NK cells. **(H)** CD11b expression on tumor-infiltrating NK cells. **(I-L)** Flow cytometry quantification of CD8^+^ T cells **(I)**, CD4^+^ T cells **(J)**, monocytes **(K)**, and macrophages **(L)** from tumor-draining lymph node of MC38 tumors from wildtype and CCR2-DTR mice. Each dot in dot plot represents a biological replicate. n.s. not significant, *p<0.05, **p<0.01, according to two-sided multiple t test with Bonferroni correction.

We also observed a higher number of NK cells in CCR2-DTR mice (Fig. 3F-G), with a similar fraction of cells in both groups expressing the maturation marker CD11b (Fig. 3H).

We also performed flow cytometry-based analysis of the tumor-draining lymph nodes, which is a critical site for T cell priming. The analysis showed increased number of CD8^+^ T cells (Fig. 3I), and a trend towards increased CD4^+^ T cells (Fig. 3J). Additionally, we observed reduced numbers of monocytes (Fig. 3K) and macrophages (Fig. 3L) in the tumor-draining lymph nodes of CCR2-depleted mice.

### CCR2 depletion enhances CD8^+^ T cell activity and synergizes with checkpoint inhibitor blockade

To investigate whether CCR2 depletion induces systemic immune changes, we performed serum cytokine profiling using the LegendPlex assay. Of the 13 cytokines assessed, six were consistently detected across samples (Supp. Fig. 3F). CCR2-DTR mice exhibited elevated levels of CCL2, indicating a compensatory response to CCR2 pathway disruption (Fig. 4A). Notably, there was also a trend of increased serum IFNγ levels, suggesting enhanced cytotoxic activities (Fig. 4B).

**Figure 4.**
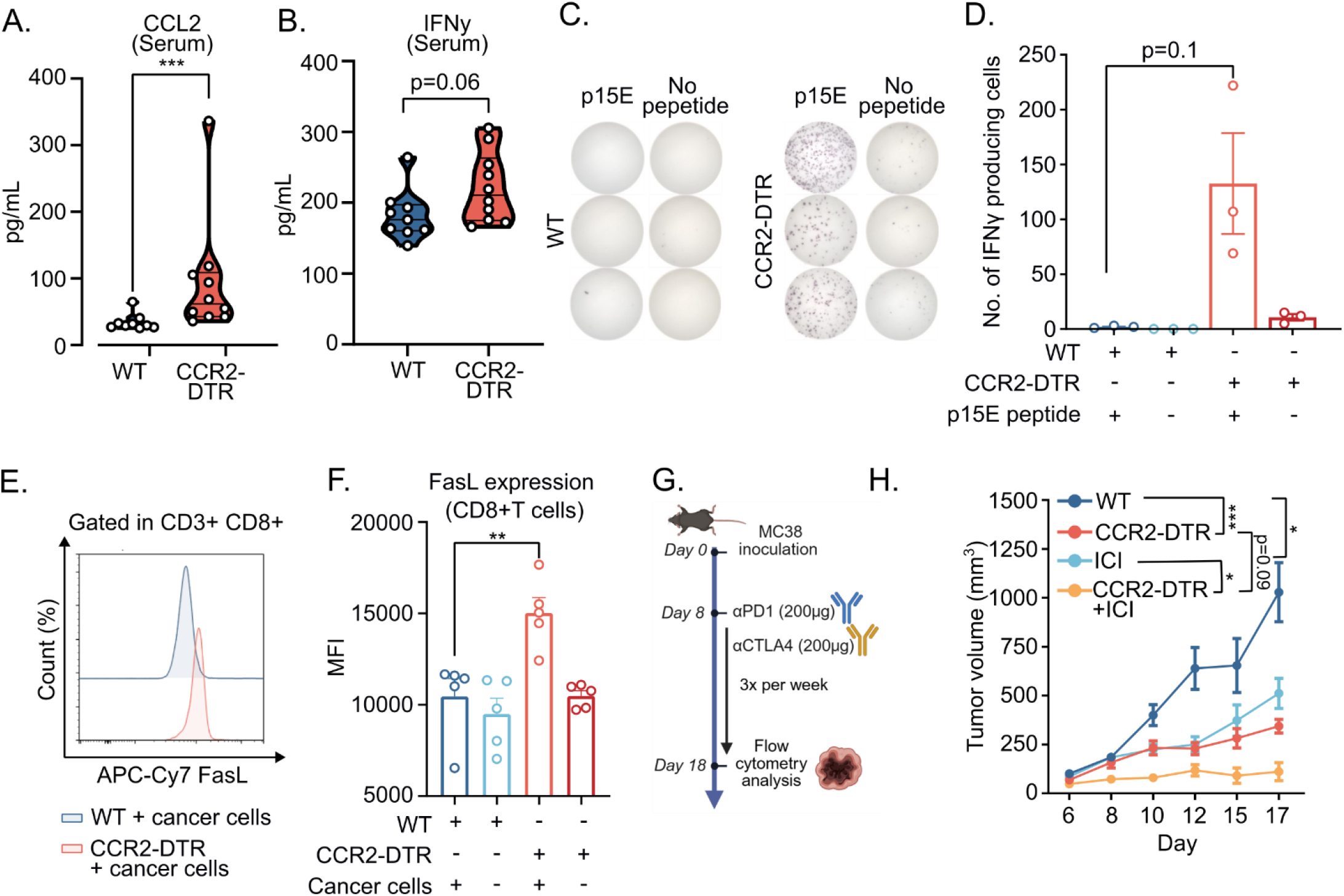
CCR2 depletion promotes CD8^+^ T cell cytotoxicity and synergizes with checkpoint inhibitor blockade. **(A-B)** Proteomics analysis using LegendPlex Mouse Inflammation (13-plex) panel was performed on serum samples collected from MC38 tumor-bearing wildtype and CCR2-DTR mice (n=10). Absolute quantity of CCL2 **(A)**, and IFNγ (**B**) in serum. **(C-D)** ELISpot was performed using 6×10^5^ splenocytes per mouse from MC38 tumor-bearing wildtype and CCR2-DTR mice (n=3 per group). Representative image **(C)** and quantification **(D)** of IFNγ ELISpot responses against p15E peptide. **(E-F)** CD8^+^ T cells were isolated from spleens of MC38 tumor-bearing wildtype and CCD2-DTR mice, and stimulated ex vivo using MC38 cells. Representative flow plot **(E)** and quantification **(F)** of FasL expression on CD8^+^ T cells. Experimental setup **(G)** and tumor growth **(H)** of combining anti-PD1 and anti-CTLA4 with CCR2^+^ cell depletion in MC38 tumor model (n=10). Each dot in dot plot represents a biological replicate. n.s. not significant, *p<0.05, **p<0.01, ***p<0.005, according to two-sided multiple t test with Bonferroni correction, or Tum Growth software for tumor growth analyses.

To further assess cytotoxic T cell activity, we isolated CD8^+^ T cells from the spleens of MC38 tumor-bearing WT and CCR2-DTR mice and stimulated them *ex vivo* with the tumor-associated antigen peptide p15E, which is highly expressed by MC38 cells (36-37). ELISpot analysis demonstrated that CD8^+^ T cells from CCR2-DTR mice generated a trend toward a higher number of IFNγ-secreting cells (Fig. 4C-D), indicating an expanded pool of tumor antigen–specific CD8^+^ cells. CD8^+^ T cells isolated from spleens of MC38 tumor-bearing CCR2-DTR mice also expressed higher levels of FasL, supporting enhanced cancer cell-specific cytotoxic activity (Fig. 4E-F).

We next evaluated whether CCR2 depletion enhances responses to immune checkpoint blockade by combining CCR2 depletion with anti–PD1 and anti–CTLA4 therapy (Fig. 4G). This combination generated a synergistic effect, resulting in delayed tumor growth (Fig. 4H) compared to either CCR2 depletion or checkpoint inhibition alone.

### CCR2 depletion promotes compensatory immunosuppressive neutrophil infiltration

Flow cytometry-based profiling of the tumor immune microenvironment revealed increased neutrophil infiltration in CCR2-depleted mice in the absence or presence of immune checkpoint blockade (Fig. 5A). We next quantified tumor-infiltrating neutrophils across multiple tumor models following CCR2 depletion and observed a similar increase in neutrophils in the MCA205 and LL2 tumor models (Supp. Fig. 4A-B). These findings indicate that neutrophil infiltration represents a conserved compensatory response to CCR2 targeting.

**Figure 5.**
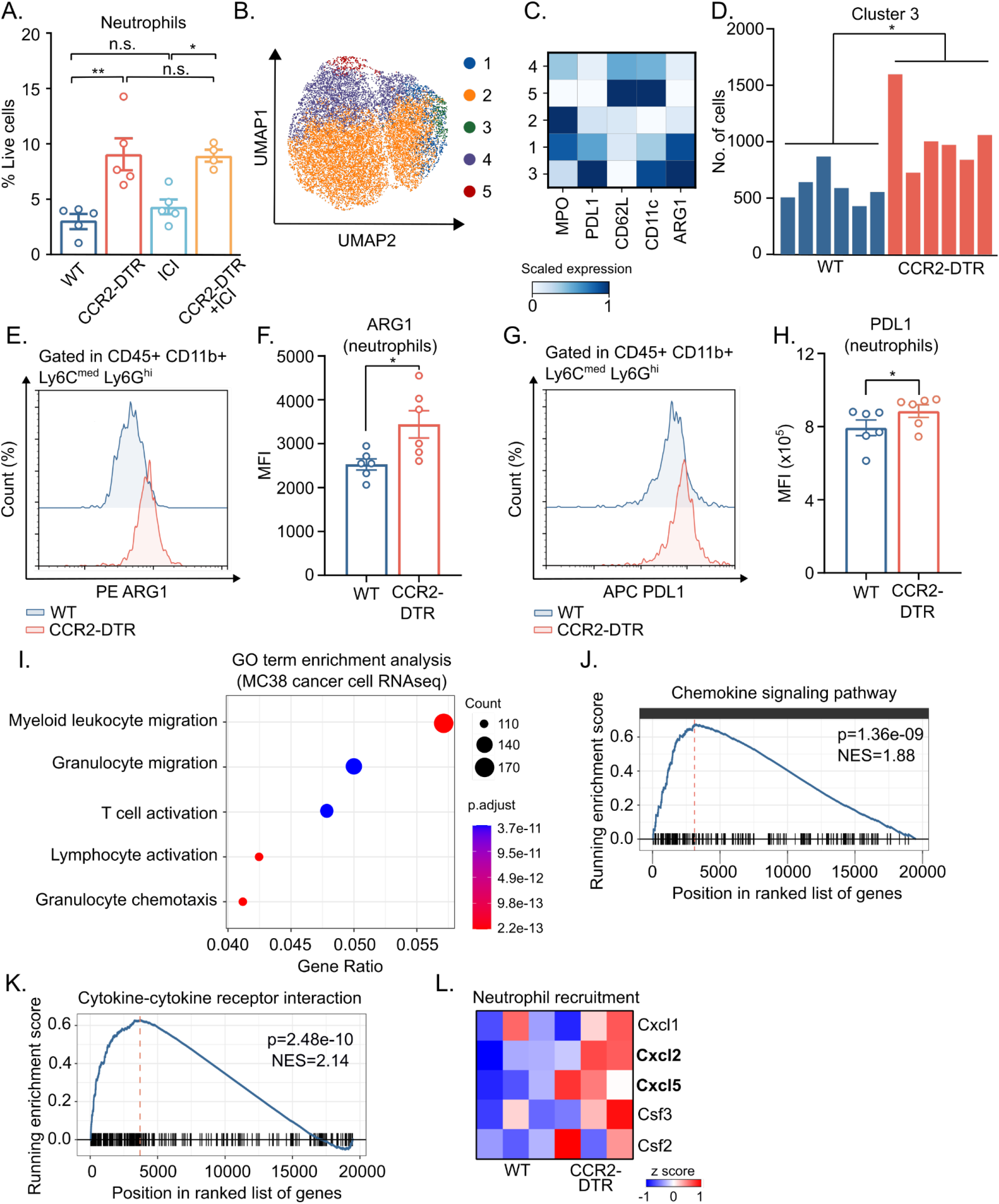
Compensatory neutrophil infiltration limits responses to CCR2 targeting. **(A-H)** Single cell suspensions were made from MC38 tumors at the experimental endpoint and analyzed by flow cytometry. **(A)** Quantification of tumor-infiltrating neutrophils. **(B-D)** MC38 tumor-infiltrating neutrophils were stained for high-dimensional flow cytometry clustering (n = 6 per group). **(B)** UMAP plots showing the distribution of FlowSOM clusters. **(C)** Heatmap of relative expression (column scaled) of markers in neutrophil clusters. **(D)** Absolute cell number in FlowSOM cluster 3. Each bar represents one biological replicate. Representative flow plot **(E)** and quantification **(F)** of ARG1 expression on tumor-infiltrating neutrophils from wildtype and CCR2-DTR mice. Representative flow plot **(G)** and quantification **(H)** of PDL1 expression on tumor-infiltrating neutrophils from wildtype and CCR2-DTR mice. **(I-L)** RNAseq was performed on MC38-GFP cancer cell isolated from tumors in wildtype and CCR2-DTR mice (n=3). **(I)** Gene ontology (GO) term enrichment analysis of upregulated genes in CCR2-DTR mice, compared to wildtype controls. **(J-K)** Plot of Gene Set Enrichment Analysis (GSEA) of KEGG pathways, showing enriched gene set in CCR2-DTR mice. **(L)** Heatmap of neutrophil recruitment molecule associated gene expression. Bold gene names indicate significantly regulated genes. Each dot in dot plot represents a biological replicate. n.s. not significant, *p<0.05, **p<0.01, according to two-sided multiple t test with Bonferroni correction.

To investigate whether the recruited neutrophils are immunosuppressive, we performed FlowSOM clustering and identified five distinct neutrophil populations (Fig. 5B-C, Supp. Fig. 4D). Cluster 1 and 2 expressed high levels of myeloperoxidase (MPO), indicating a degranulation-associated phenotype (Supp. Fig. 4E-F). Clusters 4 and 5 displayed high CD62L expression, indicating a naïve phenotype, and were more abundant in tumors from wildtype mice (Supp. Fig. 4G-H). In contrast, cluster 3 exhibited low CD62L expression, indicative of terminal differentiation, and expressed elevated levels of immunosuppressive molecules including ARG1, PDL1, and MPO. This population was significantly expanded in CCR2-DTR mice (Fig. 5D), suggesting that neutrophil infiltration may represent a compensatory mechanism that reinforces immunosuppression within the tumor microenvironment. We next quantified the expression of ARG1, PDL1, and MPO on tumor-infiltrating neutrophils from wild-type and CCR2-DTR mice. Neutrophils from CCR2-DTR tumors exhibited significantly higher ARG1 and PDL1 expression (Fig. 5E-H), with no change in MPO levels (Supp. Fig. 4I), indicating an enhanced immunosuppressive phenotype.

Since cancer cells are key sources of neutrophil recruitment chemokines (38), we performed RNAseq of cancer cells to evaluate the transcriptomic changes in cancer cells after CCR2^+^ cell depletion. RNAseq analyses showed distinct transcriptomic profiles between MC38 cancer cells isolated from tumors in wildtype and CCR2-DTR mice (Supp. Fig. 5A-C). Gene ontology analysis revealed that the top 5 upregulated pathways were predominantly immune-related (Supp. Fig. 5D). Notably, granulocyte chemotaxis and migration-related pathways as well as T cell activation-related pathways were also significantly regulated (Fig. 5I). Gene set enrichment analysis of KEGG pathways showed an upregulation in chemokine signaling pathway and cytokine-cytokine receptor interaction (Fig. 5J-K). Analysis of established neutrophil recruitment factors revealed a significant upregulation of *Cxcl2* and *Cxcl5* in cancer cells derived from CCR2-DTR mice (Fig. 5L), suggesting that tumor-intrinsic chemokine production may drive the enhanced neutrophil infiltration observed following CCR2 depletion.

### Dual CCR2 and neutrophil targeting enhances tumor control

We next investigated whether neutrophil targeting could enhance the anti-tumor effects of CCR2 depletion. Neutrophils were depleted using an antibody-mediated strategy (Fig. 6A). Due to the rapid turnover of neutrophils, we employed a dual-antibody approach combining anti-Ly6G with anti-rat κ light chain (39), and confirmed effective depletion by intracellular Ly6G staining to avoid epitope masking by the depleting antibody (Fig. 6B). CCR2^+^ cells and neutrophils were transiently depleted beginning 6 days after tumor inoculation and maintained for an additional 12 days to improve clinical translatability.

**Figure 6.**
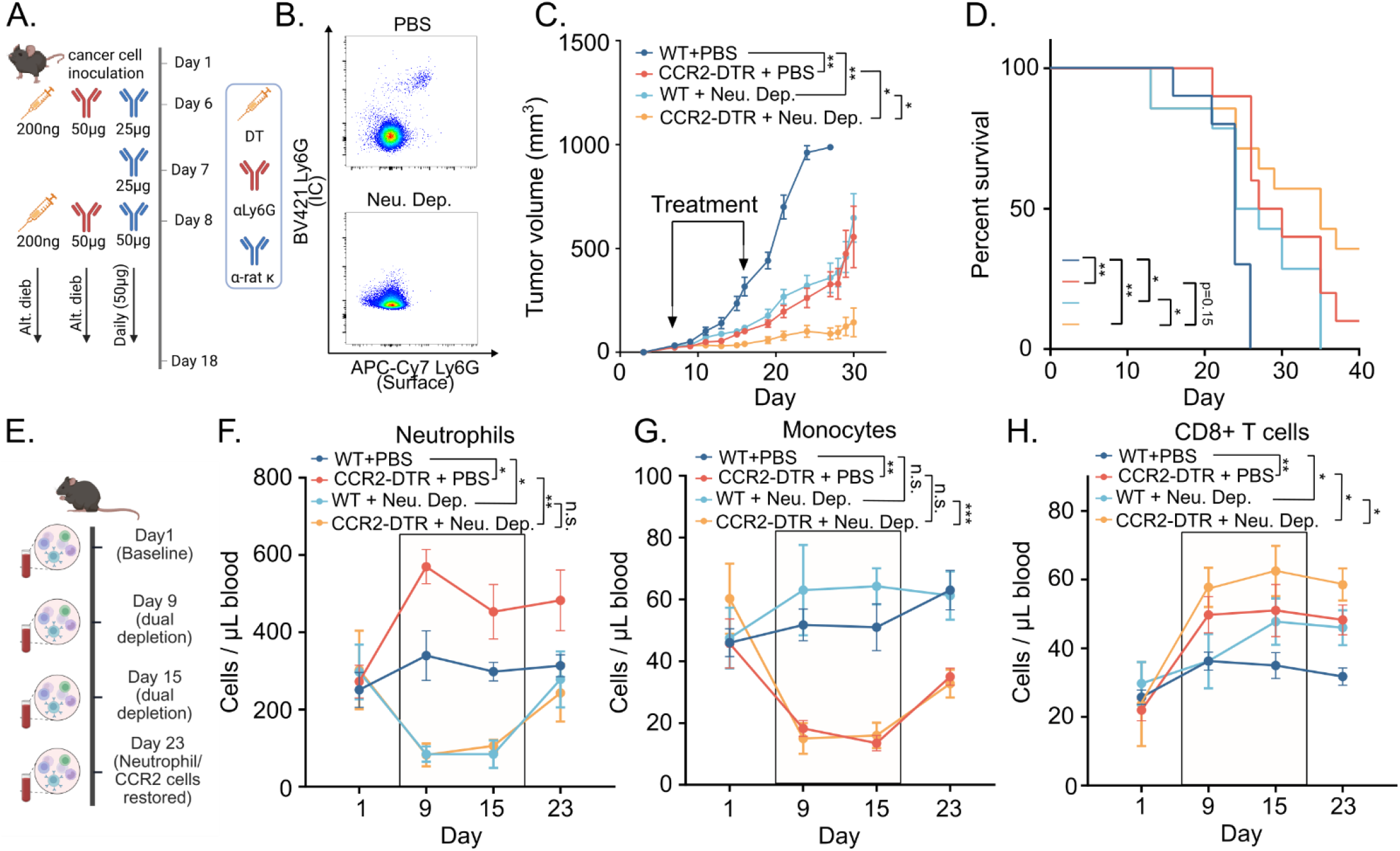
Dual CCR2 and neutrophil targeting improves tumor control. **(A)** Treatment regimen of dual CCR2^+^ cell and neutrophil targeting experiment. **(B)** Representative flow plot of surface, and intracellular (IC) Ly6G staining after neutrophil depletion (Neu. Dep.). Tumor growth **(C)** and Kaplan Meier survival plot **(D)** of dual targeting experiment. **(E)** Blood sampling schedule for peripheral immune cell monitoring. Absolute cell number of neutrophils **(F)**, monocytes **(G)** and CD8^+^ T cells **(H)** in the peripheral blood. Shaded box indicates the treatment window of dual CCR2 and neutrophil depletion. n.s. not significant, *p<0.05, **p<0.01, ***p<0.005, according to two-way ANOVA for longitudinal blood sampling analyses, or Tum Growth software for tumor growth and survival analyses.

Dual depletion markedly delayed tumor progression and prolonged survival in MC38-bearing mice (Fig. 6C-D, Supp. Fig. 6A). Complete tumor clearance was achieved in 3/12 animals, while none of the tumors regressed in the CCR2-DTR, or neutrophil depletion group (Supp. Fig. 6A). In the less immunogenic LL2 tumor model (40), we did not observe any therapeutic effect of neutrophil depletion alone; However, a small additive effect of dual depletion compared to CCR2 depletion alone was observed (Supp. Fig. 6B-C). Following tumor inoculation, we monitored peripheral circulating immune cells by flow cytometry before, during and after CCR2 depletion and /or neutrophil depletion (Fig. 6E). Similarly to the effects of CCR2 depletion on the tumor-infiltrating neutrophils, we found that CCR2 depletion also significantly increased the number of circulating neutrophils (Fig. 6F). This increase was effectively eliminated by anti-Ly6G treatment (Fig. 6F). Conversely, neutrophil depletion resulted in a trend towards elevated monocyte frequencies, indicating a reciprocal compensatory response similar to that observed with CCR2 targeting (Fig. 6G). The depletion of either CCR2^+^ cells or neutrophils led to an increase in circulating CD8^+^ T cells (Fig. 6H). The circulating CD8^+^ T cells were further expanded following transient dual depletion and persisted after treatment cessation, consistent with improved anti-tumor immune control (Fig. 6H).

## DISCUSSION

Monocytes and TAMs constitute a large fraction of immune cells in the TME (41). Inflammatory monocytes express high levels of CCR2, and are attracted to the TME by CCL2. Upon tumor infiltration they can differentiate into TAMs. Elevated levels of CCL2 are frequently observed in advanced disease stages, further exacerbating the recruitment of monocytes (13,15). Targeting immunosuppressive monocytes and TAMs by blocking the CCR2-CCL2 recruitment pathway is a promising strategy that is currently being explored in clinical settings (42,43). CCR2^+^ monocytes are highly immunosuppressive and frequently described interchangeably as CCR2^+^ monocytic myeloid-derived suppressor cells (MDSCs) (29,44). However, since there are no definite monocytic MDSC markers in the murine context (45), we here refrain from referring to CCR2^+^ cells as MDSCs without extensive ex vivo functional assays, and instead refer to them as tumor-infiltrating monocytes.

In this study, we investigated the impact of monocytes and TAMs on cancer progression and anti-tumor immunity using transgenic mice enabling DT-mediated depletion of CCR2^+^ cells. Depletion of CCR2^+^ cells reduced the numbers of tumor-infiltrating monocytes and TAMs, and resulted in significantly improved tumor control in murine models of colorectal cancer, fibrosarcoma, and melanoma. This is in line with previous studies, which found that *Ccr2* knockout mice showed reduced levels of tumor-infiltrating immunosuppressive myeloid cells, leading to reduced tumor growth and metastasis (46-49).

Although DT-mediated depletion of CCR2^+^ cells was efficient, a distinct macrophage population persisted in CCR2-DTR mice. These TAMs were characterized by the expression of the chemokine receptor CCR3. CCR3 primarily binds the eotaxins CCL11, CCL24, and CCL26, and is classically expressed on eosinophils, basophils, and T helper 2 (Th2) cells (50-53). However, under inflammatory conditions, CCR3 expression has been reported on subsets of macrophages (54). The functional roles of CCR3-expressing macrophages are not well-understood, but a previous study has shown that CCR3^+^ macrophages are required to stimulate allergen-specific T cell responses in airway inflammation (55). CCR3^+^ monocyte-derived macrophages can also drive tissue inflammation in non-alcoholic fatty liver disease, where their transcriptional programs are enriched for inflammatory cytokine secretion (56). In the context of cancer, our transcriptomic analyses demonstrated that CCR3^+^ TAMs in CCR2-DTR tumors displayed a pro-inflammatory phenotype, characterized by reduced gene expression of immunosuppressive molecules *Il10*, and *Ptgs2*. These findings indicate that CCR2 depletion spares a subset of pro-inflammatory CCR3^+^ TAMs, which might have important anti-tumorigenic functions.

In addition to remodeling the TAM compartment, CCR2 targeting exerted broad effects on the lymphocyte compartment. We demonstrated that CCR2 depletion led to an increased intratumoral infiltration of CD8^+^ T cells with elevated PD1 expression. Previous studies have indeed shown that CCR2^+^ myeloid cells can restrict antigen-specific CD8^+^ T cell trafficking, and directly suppress T cell function through contact-dependent mechanisms or ARG1- and iNOS-mediated mechanisms (44). CCR2^+^ cells are also recruited to the TDLN and impair T cell priming (57). In line with this, we showed that CCR2 depletion increased the number of CD8^+^ T cells in the TDLN. CCR2 depletion also expanded the pool of tumor antigen–specific effector T cells, as reflected by increased numbers of IFNγ-producing splenocytes and elevated FasL expression on CD8^+^ T cells. This was accompanied by a trend toward higher systemic IFNγ levels in the serum. Improved CD8+ T cell-mediated anti-tumor immunity upon CCR2 targeting has been shown in previous studies in murine models of lung cancer, breast cancer, melanoma lung metastasis, and pancreatic cancer, using CCR2 knockout mice or CCR2/CCL2 blockade (27,28,58). In this study, we found that CCR2 depletion synergized with checkpoint inhibitors in controlling tumor growth. This is consistent with previous studies showing synergy between CCR2 inhibition and checkpoint inhibitors in other tumor models such as glioma and lung cancer (27,30). These findings indicate that CCR2 targeting restores cytotoxic T cell function and enhances anti-tumor immunity, providing a mechanistic basis for its synergy with checkpoint inhibitors.

Despite the efficacy of combining CCR2 depletion with checkpoint inhibitors, a subset of mice remained resistant to treatment. One possible explanation is the activation of compensatory responses within the myeloid compartment, which is highly plastic and adapts rapidly to therapeutic intervention (59,60). Through systematic profiling of the intratumoral myeloid landscape following CCR2 depletion, we identified neutrophil infiltration as a key compensatory mechanism that limits therapeutic efficacy. CCR2 targeting consistently increased neutrophil infiltration into the TME across multiple tumor models. Tumor-infiltrating neutrophils (TANs) have emerged as critical regulators of anti-tumor immunity, and have been shown to promote tumor progression through diverse mechanisms, such as direct suppression of T cells, facilitating metastatic dissemination, and support of tumor angiogenesis (61,62). In our study, flow cytometry analysis revealed that TANs induced by CCR2 depletion are terminally differentiated and express high levels of immunosuppressive mediators ARG1 and PDL1, indicating their capacity to inhibit T cell function. We identified cancer cells as a source of neutrophil recruitment chemokines *Cxcl2* and *Cxcl5* in the TME upon CCR2^+^ cell depletion through RNA sequencing. Similar neutrophil-driven resistance mechanisms have been reported upon TAM-targeting CSF1R blockade, where compensatory neutrophil infiltration undermines therapeutic benefit across multiple murine tumor models, such as glioblastoma, breast, lung, and melanoma (63,64). CSF1R inhibition promotes the recruitment of neutrophils capable of potently suppressing antigen-driven splenocyte proliferation (63).

When combined with antibody-mediated neutrophil depletion, CCR2 targeting showed strong synergy, resulting in marked tumor control, prolonged survival, and complete tumor clearance in 25% of treated mice. Consistent with our observations, prior studies have shown that combining macrophage-targeting CSF1R blockade with the inhibition of CXCR2, a key chemokine receptor mediating neutrophil recruitment, significantly improves anti-tumor responses (63). Several neutrophil-targeting strategies are currently under development, including approaches that limit neutrophil recruitment via CXCR2/CXCR1 blockade or target neutrophil-induced immunosuppression (65,66), opening possibilities for novel combination strategies. In addition to targeted agents, certain chemotherapy regimens may indirectly modulate neutrophil dynamics. Notably, a small phase Ib study combining CCR2 blockade with FOLFIRINOX chemotherapy reported preliminary evidence of clinical activity (67). In pancreatic cancer patients, severe neutropenia following the FOLFIRINOX regimen has been associated with improved overall survival (68), suggesting that part of its efficacy may derive from effects on immunosuppressive neutrophils. Together, these observations suggest that chemotherapy-mediated neutrophil modulation could also synergize with CCR2 inhibition.

Collectively, our findings highlight the therapeutic potential of CCR2 targeting, provide a mechanistic rationale for combination with immune checkpoint blockade, and support further evaluation of combination strategies incorporating neutrophil-targeting agents or neutrophil-modulating chemotherapies to overcome resistance and maximize clinical benefit in patients.

## MATERIALS AND METHODS

### Cell lines

MC38, MC38-GFP, B16F10 (B16), LL2, T241-tdTomato (T241) cell lines were retrieved from CCIT-DK cell line biobank. MC38, B16, and LL2 cells were cultured in growth media consisting of DMEM (10564011), 10% FBS (A5209501), 1% P/S (15140122), and 1% HEPES (15630080) (All from Gibco). T241 cells were cultured in growth media consisting of RPMI 1640 (61870036), 15% FBS (A5209501), 1% HEPES (15630080) and 1% P/S (15140122) (All from Gibco). MCA205 cell line was purchased from Merck (SCC173) and cultured in complete growth medium, consisting of RPMI-1640 GlutaMAX™ Supplement HEPES, 20% FBS, 1% sodium pyruvate, 1% nonessential amino acids, and 1% P/S (All from Gibco). The complete growth medium was supplemented with β-mercaptoethanol (Gibco) at a concentration of 55 µM before use.

### In vivo experiments

Animal experiments were conducted at the animal facility of the Department of Oncology, Herlev Hospital, Denmark, under licenses granted by the Danish Animal Experiments Inspectorate. All mice were housed in a 22 °C environment and subjected to a standard 12-hour dark and light cycle. All procedures were conducted in accordance with the approved licenses. CCR2-DTR mice were obtained from Dr. Eric G. Pamer (69), Memorial Sloan Kettering Cancer Center. Daily maintenance of CCR2-DTR mouse stocks was performed by the animal caretakers at the animal facility. CCR2-DTR heterozygous males were bred with wildtype C57BL/6 females, and both transgenic and wildtype littermates were used in each experiment. Genotyping of CCR2-DTR mice for the targeted allele (DTR) was performed with the primers DTR_fw (5’ to 3’): ACGAGAAGCGCGATCACAT and DTR_rv (5’ to 3’): ATCGTAGTCATACGGTGTGGTG. All experimental mice were females, 8 to 20 weeks of age, and were co-housed.

For subcutaneous tumor models, 0.5 × 10^6^ MC38, or MC38-GFP, or B16, or LL2, or T241 cells were injected subcutaneously on the right flank per mouse. For MCA205, 1 × 10^6^ cells were injected subcutaneously on the right flank per mouse. Tumor volumes were measured at least three times a week with digital calipers. Experimental endpoint was tumor volume exceeding 1000mm^3^, calculated by the formula: 0.5 x length x width^2^. Humane endpoint was weight loss more than 20%, tumor ulceration, or tumor wounds of more than 5mm. Mice were euthanized by cervical dislocation, tumors were excised and collected for further analyses. Mice euthanized due to tumor ulcerations were excluded from the analysis.

### In vivo treatments

For induction of cell depletion in CCR2-DTR mice, 8 - 10 ng/g body weight diphtheria toxin (D0564, Sigma) was injected intraperitoneally (i.p.) two days after cancer cell inoculation (unless otherwise stated in the figure) for both CCR2-DTR mice and wildtype littermates, and continued 3 times per week until experimental endpoint.

For checkpoint inhibitor experiments, mice were treated i.p. three times a week, starting 5 days after cancer cell inoculation, with αPD1 monoclonal antibody (mAb) (200 μg/mouse; BE0146, clone RMP1-14, BioXcell), or αCTLA4 mAb (200 μg/mouse; BE0164, clone 9D9, BioXcell) until endpoint. Control mice were treated with PBS according to the same schedule.

For antibody-mediated neutrophil depletion, anti-Ly6G (clone 1A8, #BP0075-1), and anti-rat Kappa immunoglobulin (clone MAR 18.5, #BE0122) were injected i.p. according to the experimental set up in the figure. Briefly, anti-Ly6G was started on day 6 after cancer cell inoculation, at a dosage of 50μg per mouse every other day. Anti-rat Kappa immunoglobulin was injected every other day, starting on day 6 at an initial dosage of 25μg per mouse for two treatments, and a follow-up dosage of 50μg per mouse until treatment cessation. When mice were sequentially injected with two antibodies, a 2-hour delay was maintained between injections. Control mice were treated with PBS according to the same schedule.

### Flow cytometry analyses

For analysis of tumor-infiltrating immune cells, mice were euthanized by cervical dislocation, and tumor excised. The tumors were minced with scissors, and enzymatically digested in RPMI medium (Gibco) containing 2.1 mg/ml collagenase type 1 (LS005275, Worthington), 75 μg/ml DNase I (LS006331, Worthington), 5 mM CaCl_2_, and 1% P/S for 1 hour at 37 °C and 300RPM in a thermo shaker (VWR). The tubes were inverted to mix at the 30-min time point. The single-cell suspension was filtered through a 70μm cell strainer (Corning), and washed with 10mL of cold FACS buffer (PBS with 0.5% bovine serum albumin (Sigma-Aldrich) and 5 mM EDTA (Sigma-Aldrich)).

For peripheral immune cell analysis, lateral tail vein of mice was punctured with a sterile 25G needle, and 30 μL blood per mouse was transferred into round bottom 96 well plate, containing 150μL of 5 mM EDTA in PBS per well. Tumor draining lymph nodes were excised and dissociated mechanically by pressing with a syringe plunger against a 70 μm cell strainer followed by a second filtration through a 70 μm cell strainer. For bone marrow cell isolation, femurs were dissected and rinsed first in 70% ethanol and subsequently in PBS, and the bone marrow cells were flushed with PBS using a sterile 25 G needle.

Prior to staining, red blood cells were lysed with RBC Lysis Solution (158904, Qiagen) at room temperature for 5 minutes, and washed with cold FACS buffer. Cells were then incubated with FcR block (130-092-575, Miltenyi) for 20 min at 4 °C. Dead cells were excluded using Zombie Aqua viability dye (423101, BioLegend). For surface staining, cells were incubated with antibodies at 4 °C in the dark for 30 min. Cells were then washed twice with FACS buffer, fixed with Perm/Fix buffer (554714, BD Biosciences) for 30 min at 4^°^C. For intracellular markers, the cells were subsequently stained with intracellular antibodies at 4^°^C in the dark for 45 min.

For cytotoxic molecule staining, CD8^+^ T cells were isolated from spleens of tumor-bearing mice using CD8a (Ly-2) MicroBeads (130-117-044, Miltenyi). Purity of isolated cells was routinely tested to be more than 90%. Isolated CD8^+^ T cells were stimulated for 20 hours with target cells, and a further 4-hour stimulation in the presence of 1x Golgi plug (BDB555029, BD Biosciences). CD8^+^ T cells were washed once with FACS buffer and stained for surface markers according to above protocol.

All data were acquired on FACSCanto II (BD Biosciences) or Novocyte Quanteon (Agilent) flow cytometer within 3 days of staining and fixation after appropriate compensation using compensation beads (01-2222-41, BD Biosciences). Manual gating was performed using FlowJo software (Tree Star) or NovoExpress software (Agilent). High dimensional flow cytometry data clustering was performed with CRUSTY web platform (https://crusty.humanitas.it/) (70) with the following parameters: runtime - full, clustering type - flowSOM, batch correction - none, UMAP distance − 0.5, UMAP spread − 1. Antibodies used for flow cytometry are listed in Supplementary Table 1.

### Enzyme-linked immune-spot (ELISpot) assay

Splenocytes from tumor-bearing wildtype and CCR2-DTR mice were isolated as described above. 96-well plates were coated overnight at 4°C with an anti-mouse IFN-γ capture antibody (12 μg/mL; 3321-3-1000, MabTech). The plates were blocked using RPMI-1640, 10%FBS, and 1%P/S for 30 min at 37°C. 6 × 10^5^ splenocytes per well were stimulated with peptide for the cancer-associated antigen p15E (KSPWFTTL, Schäfer-N) at a concentration of 5 μM overnight at 37°C. Unstimulated splenocytes were used as negative controls. The plates were developed using a biotinylated anti-mouse IFN-γ detection antibody (3321-6-250, MabTech), streptavidin-ALP (3310-10, MabTech), and BCIP/NBT substrate (3650-10, MabTech). Spots were quantified using a CTL ImmunoSpot S6 analyzer with ImmunoSpot software (v5.1, CTL Analyzers). All conditions were run in technical triplicates.

### RNA sequencing

For tumor-infiltrating macrophage isolation, single-cell suspensions of MC38 tumors from wildtype and CCR2-DTR mice were labeled with anti-F4/80 microbeads (130-110-443, Miltenyi) according to manufacturer’s instructions, and loaded onto LS columns (130-042-401, Miltenyi). The F4/80^+^ fractions were collected in MACS buffer (PBS + 2% FBS + 2 mM EDTA), and loaded onto a new LS column to enhance purity. The F4/80^+^ fraction, representing macrophages, were collected. The purity of macrophages was tested to be at least 92%.

For MC38 cancer cell isolation, single-cell suspensions of MC38-GFP tumors from wildtype and CCR2-DTR mice were labelled with Zombie Aqua (423101, BioLegend) viability dye. Live GFP^+^ cells were sorted on FACSMelody cell sorter (BD Biosciences). The purity of cancer cells was tested to be at least 96%.

Isolated macrophages and cancer cells were stored in buffer RLT at −80 °C until RNA extraction using the RNeasy Mini Kit (74104, Qiagen), according to the manufacturer’s instructions. The quality of RNA samples was determined by Bioanalyzer (Agilent). Samples with RNA integrity number (RIN) of at least 6.5 were used for RNA sequencing.

In total, 500 ng RNA from cancer cells, and 200ng RNA from macrophages per sample were prepared for sequencing using polydT enrichment according to the manufacturer’s instructions (Illumina). Library preparation was performed using the NEBNext RNA library prep kit for Illumina. The library quality was assessed using a Fragment Analyzer followed by library quantification using the Illumina library quantification kit. Libraries were sequenced on a NovaSeq 6000 platform (Illumina). Sequenced reads were aligned to the reference mm39 genome for mouse using STAR, version 2.7.9. The gene expression count matrix was generated using featureCounts version 1.6.4 and GENCODE gene annotation with the parameters “-p -t exon”. RNAseq data can be accessed on GEO repository (GSE329852).

For all analyses of the RNA seq data, R version 4.3.2 was used. Analysis of differentially expressed genes was performed using DESeq2 package version 1.42.0 (cut off p-adjusted value < 0.05 and log2 fold change >1.5). For the GO term enrichment analysis, enrichGO from the clusterProfiler package version 4.10.0 was used. The Benjamini-Hochberg method and p-adjusted-value cut-off <0.05 were set as criterion for the GOSeq analysis. The ggplot2 package version 3.3.3, was used to visualize the GOterms. For GSEA analysis of KEGG pathways, gseaKEGG from the clusterPRofiler package version 4.10.0 was used, and p-adjusted value cut-off <0.05 were set as a criterion for the analysis. The gseaplot from enrichplot package version 1.22.0 was used to visualize the regulated pathways.

For the estimation of M1 and M2 macrophages percentages, immune cell infiltration deconvolution tool quanTIseq (33) was assessed through the TIMER2.0 web platform (71), using the following settings: species - “mouse”, cancer type - “AUTO”. Estimation of M1 and M2 macrophages was downloaded from the web platform and visualized using R studio (version 4.3.2).

### Proteomics

Terminal blood samples were collected via decapitation from MC38 tumor bearing wildtype and CCR2-DTR mice. Blood samples were allowed to coagulate at room temperature for 30 minutes before centrifugation at 1200xg for 10 minutes. Serum was carefully transferred to new tubes using a pipette. Serum samples were used fresh, and undiluted. The assay was performed according to the manufacturer’s instructions using the LEGENDplex™ Mouse Inflammation Panel (13-plex) (cat. no. 740150, BioLegend). Data was acquired on Novocyte Quanteon (Agilent). The assay FCS files were analyzed using BioLegend’s LEGENDplex™ data analysis software.

## Statistical analyses

Statistical analyses were performed using two-sided multiple *t* test with Bonferroni-Dunn correction for multiple comparisons in Prism 8 (GraphPad). Longitudinal peripheral immune cell data were analyzed using two-way ANOVA in Prism 8 (GraphPad). Tumor growth-related statistics were analyzed using TumGrowth software (72). Survival data were analyzed using log-rank tests in Prism 8 (GraphPad).

## Acknowledgments

We thank animal technicians Anne Boye and Ditte Stina Jensen for daily housekeeping work at the animal facility and assistance with animal experiments.

## Author contributions

Conceptualization: KYF, MC, DHM. Data curation: KYF, AZJ, HL, MC, MS, MLH, LG. Formal analysis: KYF, AZJ, HL, MC, LG. Investigation: KYF, AZJ, HL, MC, KJB, MLH MS, M-LT. Resources: LG, DHM. Supervision: LG, MHA, DHM. Visualization: KYF, AZJ, DHM. Writing – original draft: KYF and DHM. Writing – review & editing: KYF, AZJ, HL, MC, MS, KJB, M-LT, MLH, MHA, LG, DHM.

## Data and materials availability

All data are available in the main text or the supplementary materials. RNA sequencing data have been deposited in the GEO repository under accession number GSE329852.

