## Supplementary figure 1-6, supplementary table 1 for "Dual targeting of CCR2^+^ monocytes and neutrophils enhances anti-tumor immunity"

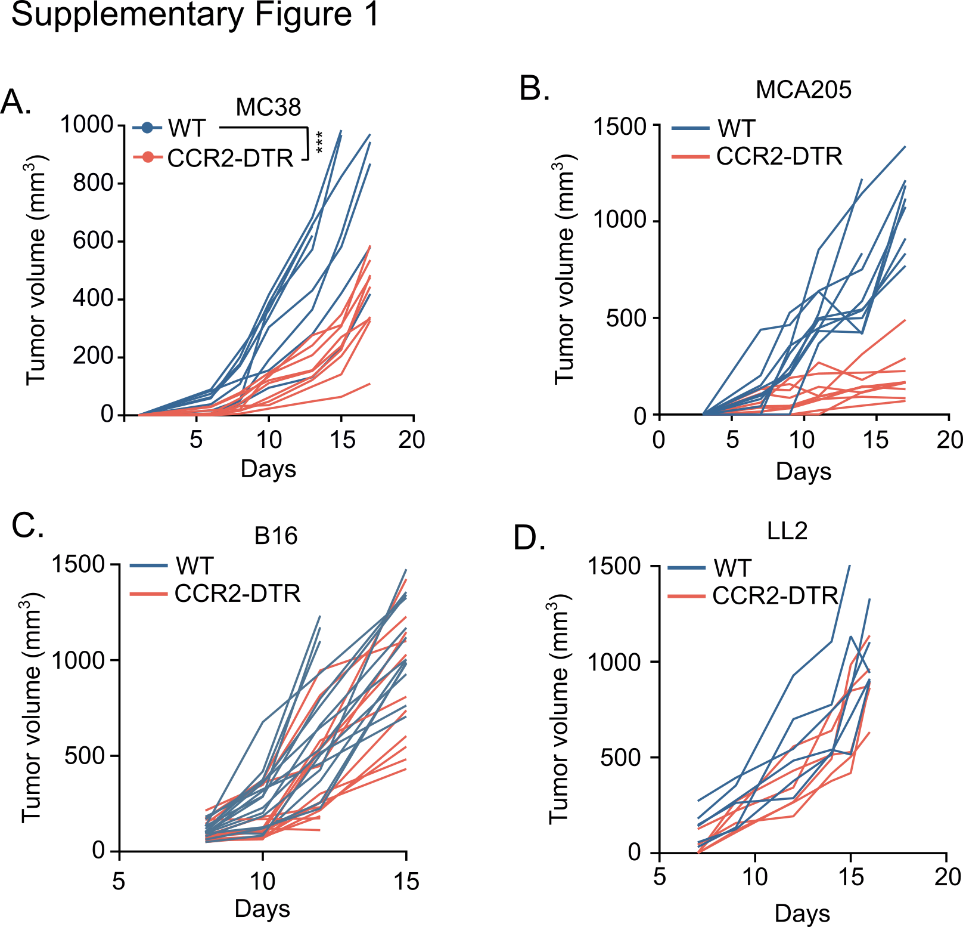

**Supplementary figure 1.**

Tumor growth kinetics of MC38 **(A)**, MCA205 **(B)**, B16 **(C)**, and LL2 **(D)** in wildtype or CCR2-DTR mice. Each line in tumor growth plot represents one biological replicate.

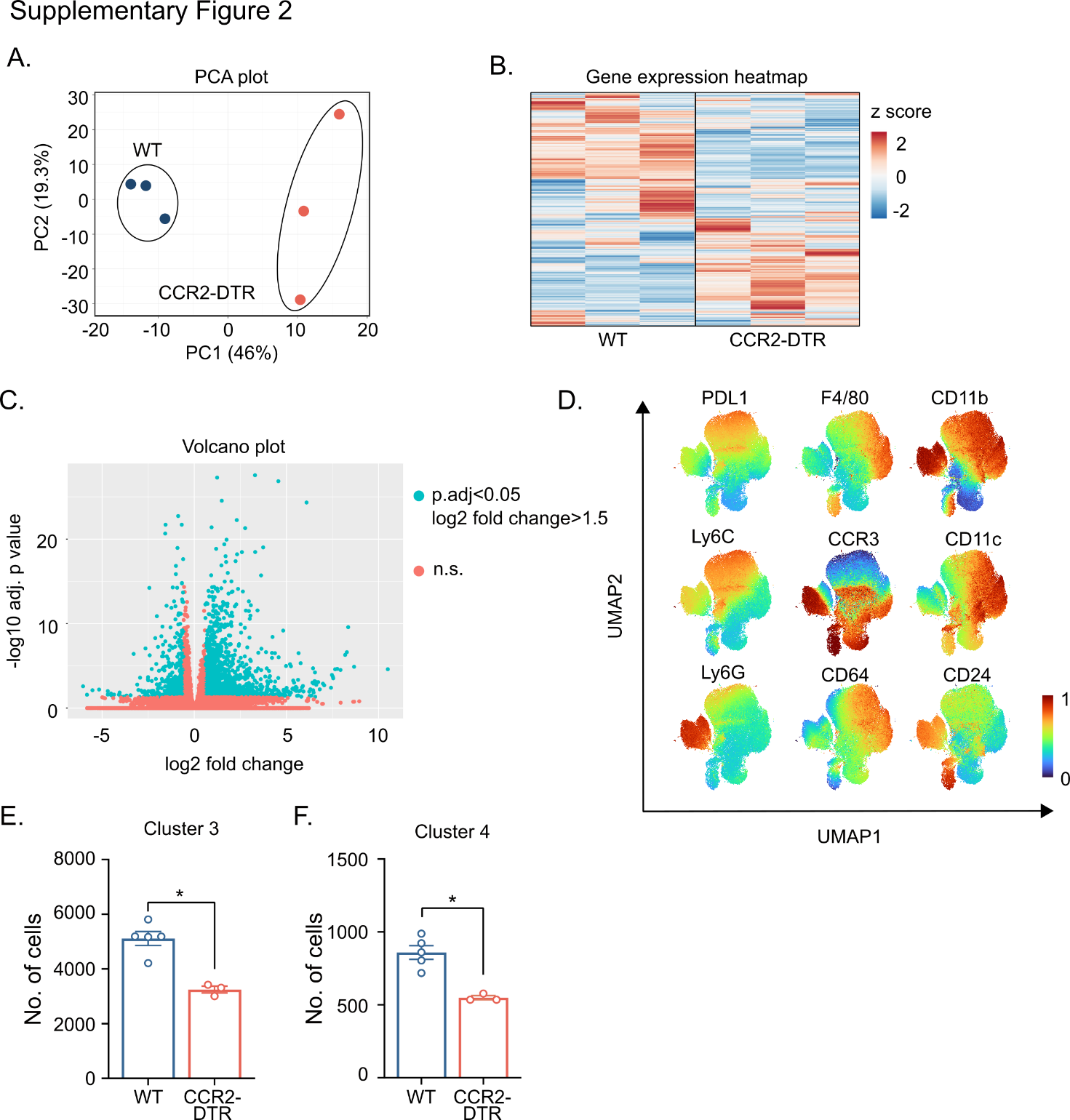

**Supplementary figure 2.**

**(A-C)** TAMs were isolated from MC38 tumors, and subjected to RNAseq. RNAseq analysis showing PCA plot **(A)**, heatmap **(B)**, and volcano plot (**C**) of genes differentially regulated in TAMs from wildtype and CCR2-DTR mice. (**D–F)** Tumor-infiltrating myeloid cells (CD11c^+^ or CD11b^+^) were stained for FlowSOM clustering (n = 3-5 per group). (**D)** Tiled UMAP plots showing relative expression (z score) of markers by tumor-infiltrating myeloid cells. **(E-F)** Dot plots showing percentages of FlowSOM cluster 3 **(E)** and 4 **(F)**. Each dot in dot plots represents one biological replicate. *p<0.05, according to two-sided multiple t test with Bonferroni correction. Mean ± SEM are depicted.

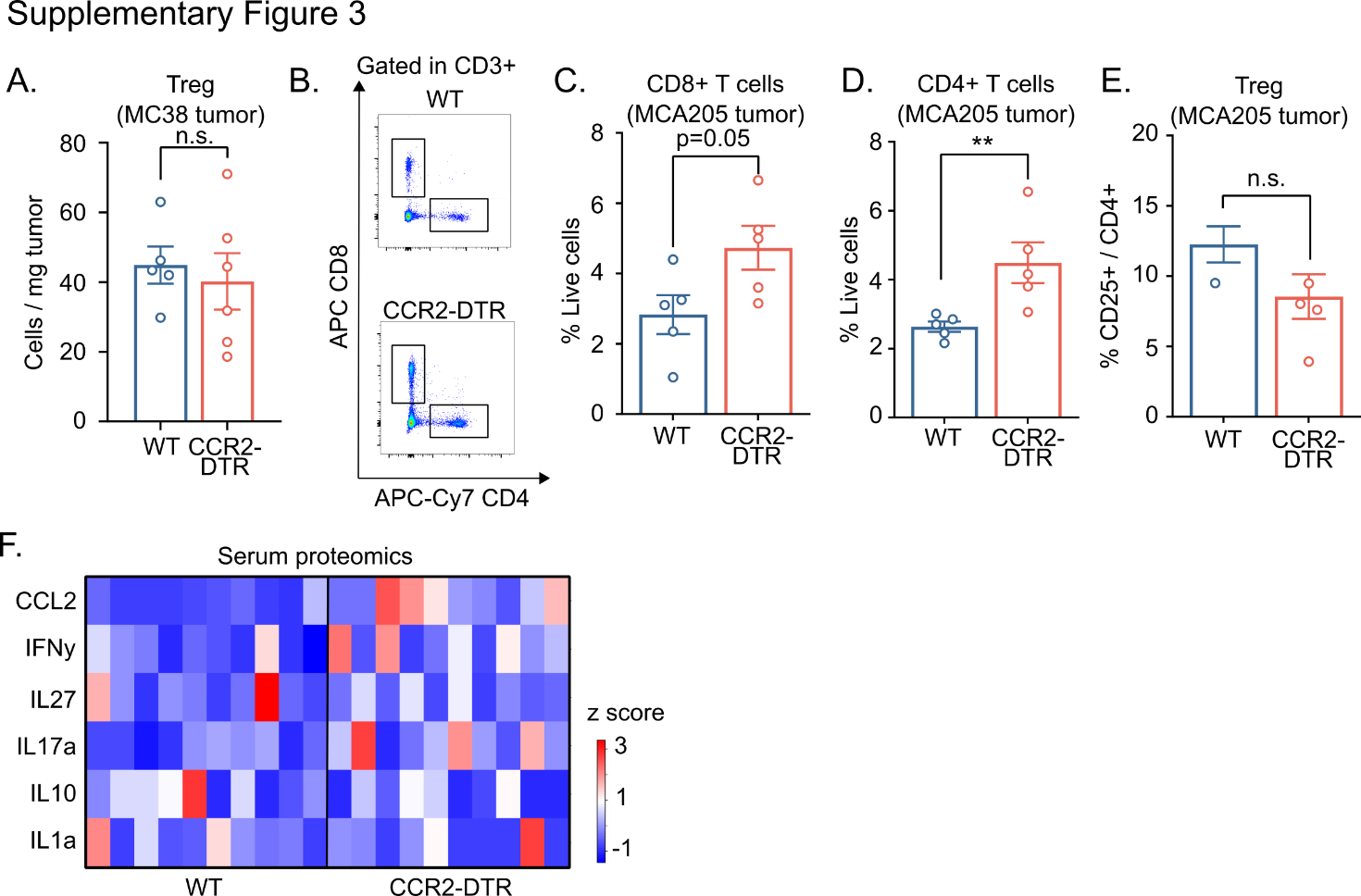

**Supplementary figure 3.**

Single cell suspensions were made from MC38 **(A)**, MCA205 **(B-G)**, and LL2 **(H)** tumors in wildtype and CCR2-DTR mice at the experimental endpoint and analyzed by flow cytometry. **(A)** Quantification of Tregs in MC38 tumors. Representative flow plot **(B)** and quantification of CD8+ T cells **(C)** and CD4+ T cells **(D)** in MCA205 tumors. **(E)** Quantification of Treg in MCA205 tumors. **(F)** Proteomics analysis using LegendPlex Mouse Inflammation (13-plex) panel was performed on serum samples collected from MC38 tumor-bearing wildtype and CCR2-DTR mice (n=10). **(F)** Heatmap of relative quantity (z score) of serum protein markers detected. Each dot in dot plots represents one biological replicate. n.s. not significant, **p<0.01, according to two-sided multiple t test with Bonferroni correction. Mean ± SEM are depicted.

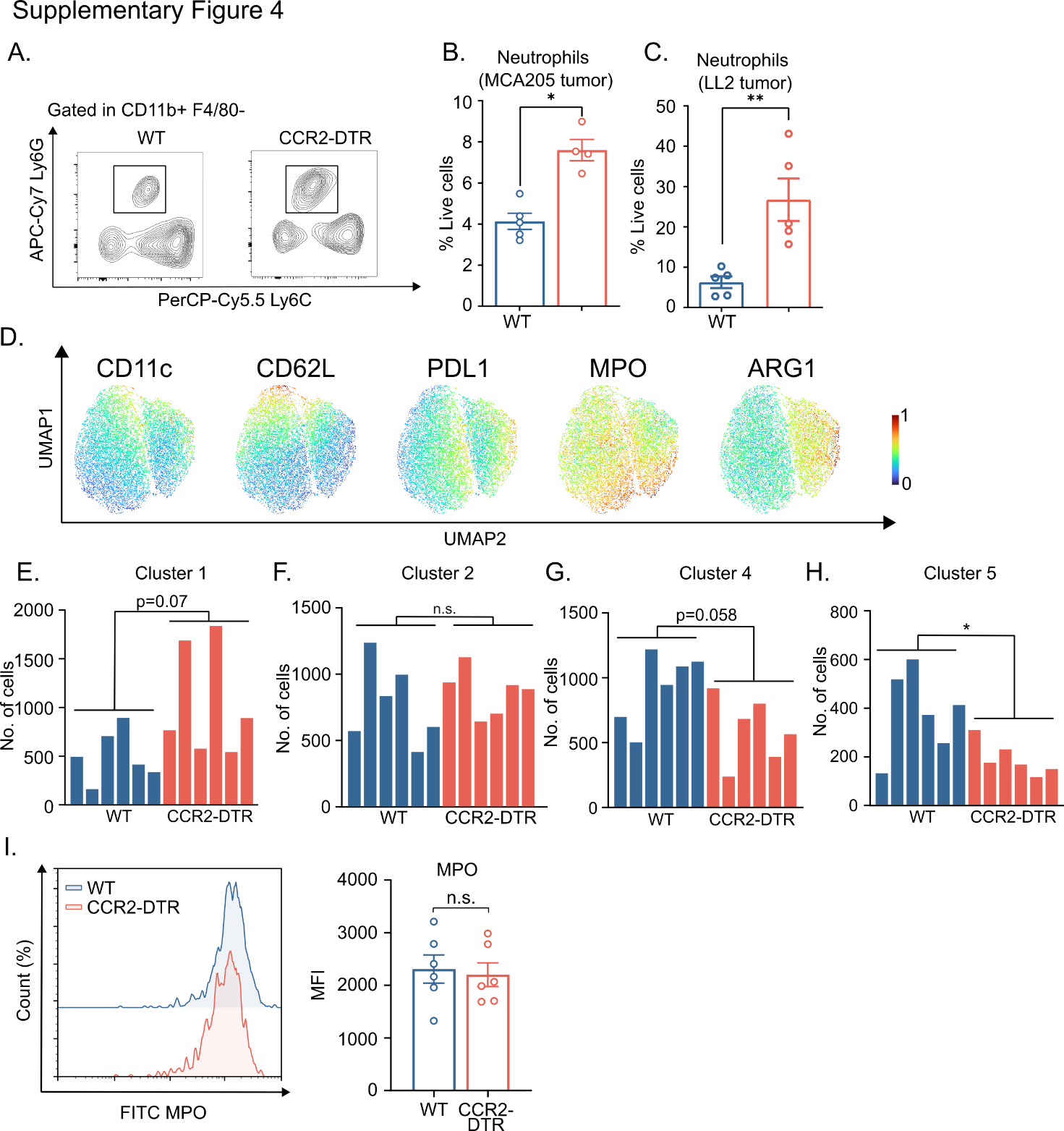

**Supplementary figure 4.**

Representative flow plot **(A)** and quantification **(B)** of tumor-infiltrating neutrophils in MCA205 tumors. **(C)** Quantification of tumor-infiltrating neutrophils in LL2 tumors. **(D-H)** Tumor-infiltrating neutrophils (CD11b^+^ F4/80^-^ Ly6G^+^ Ly6Cmed) were stained for FlowSOM clustering (n = 6 per group).  (**D)** Tiled UMAP plots showing relative expression (z score) of markers by tumor-infiltrating neutrophils. **(E-H)** Bar plots showing percentages of FlowSOM cluster 1 **(E)**, cluster 2 **(F)**, and cluster 4 **(G)**, and cluster 5 **(H)**. **(I)** Representative histogram (left) and quantification (right) of MPO expression in tumor-infiltrating neutrophils. Each dot in dot plots represents one biological replicate. n.s. not significant, *p<0.05, **p<0.01 according to two-sided multiple t test with Bonferroni correction. Mean ± SEM are depicted.

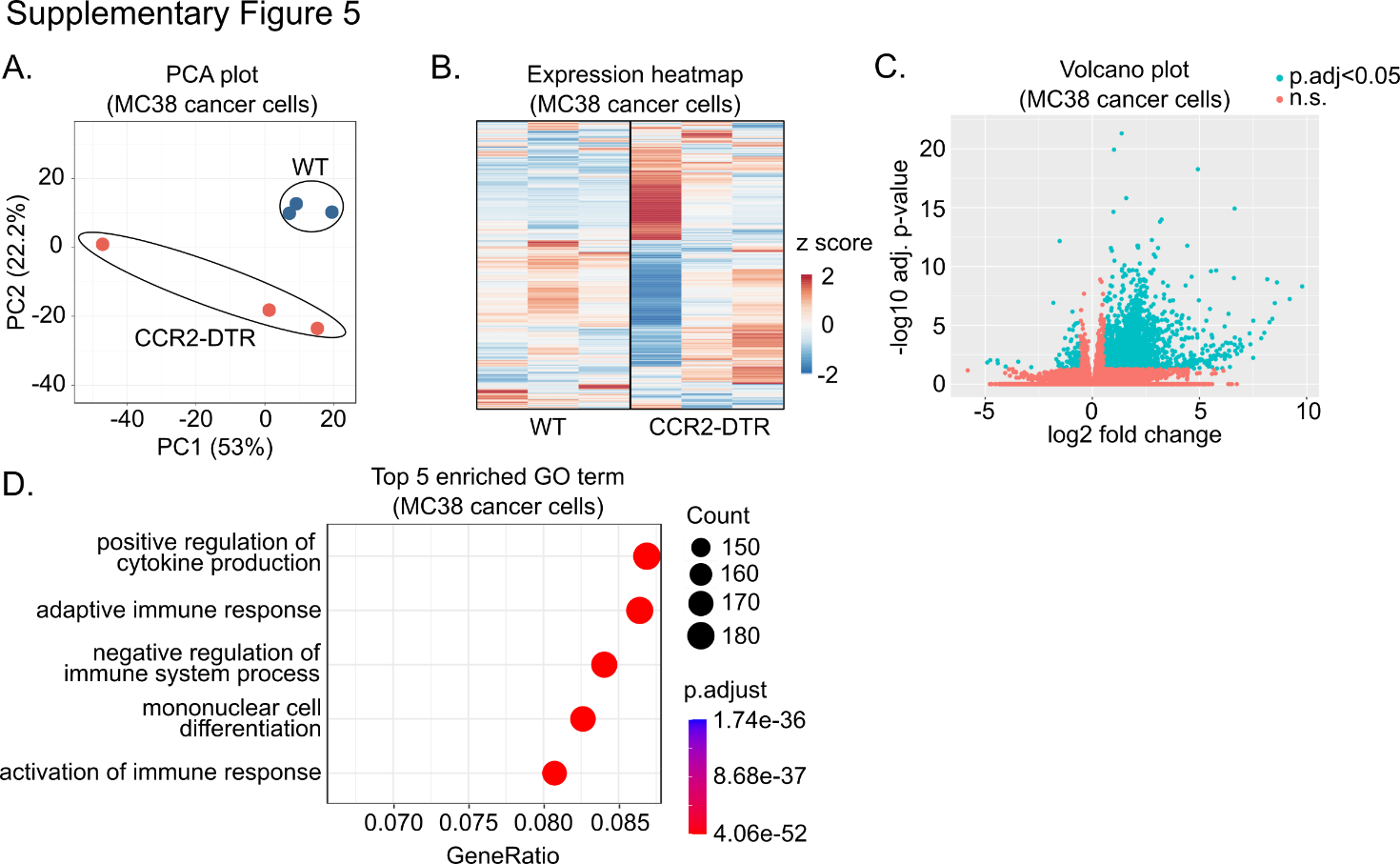

**Supplementary figure 5.**

**(A-D)** RNAseq was performed on MC38-GFP cancer cell isolated from tumors in wildtype and CCR2-DTR mice (n=3). RNA-seq analysis showing PCA plot **(A)**, heatmap **(B)**, and volcano plot (**C**) of genes differentially regulated in cancer cells from wildtype and CCR2-DTR mice. **(D)** Gene ontology (GO) term enrichment analysis of upregulated genes in CCR2-DTR mice, compared to wildtype controls, showing top 5 enriched terms.

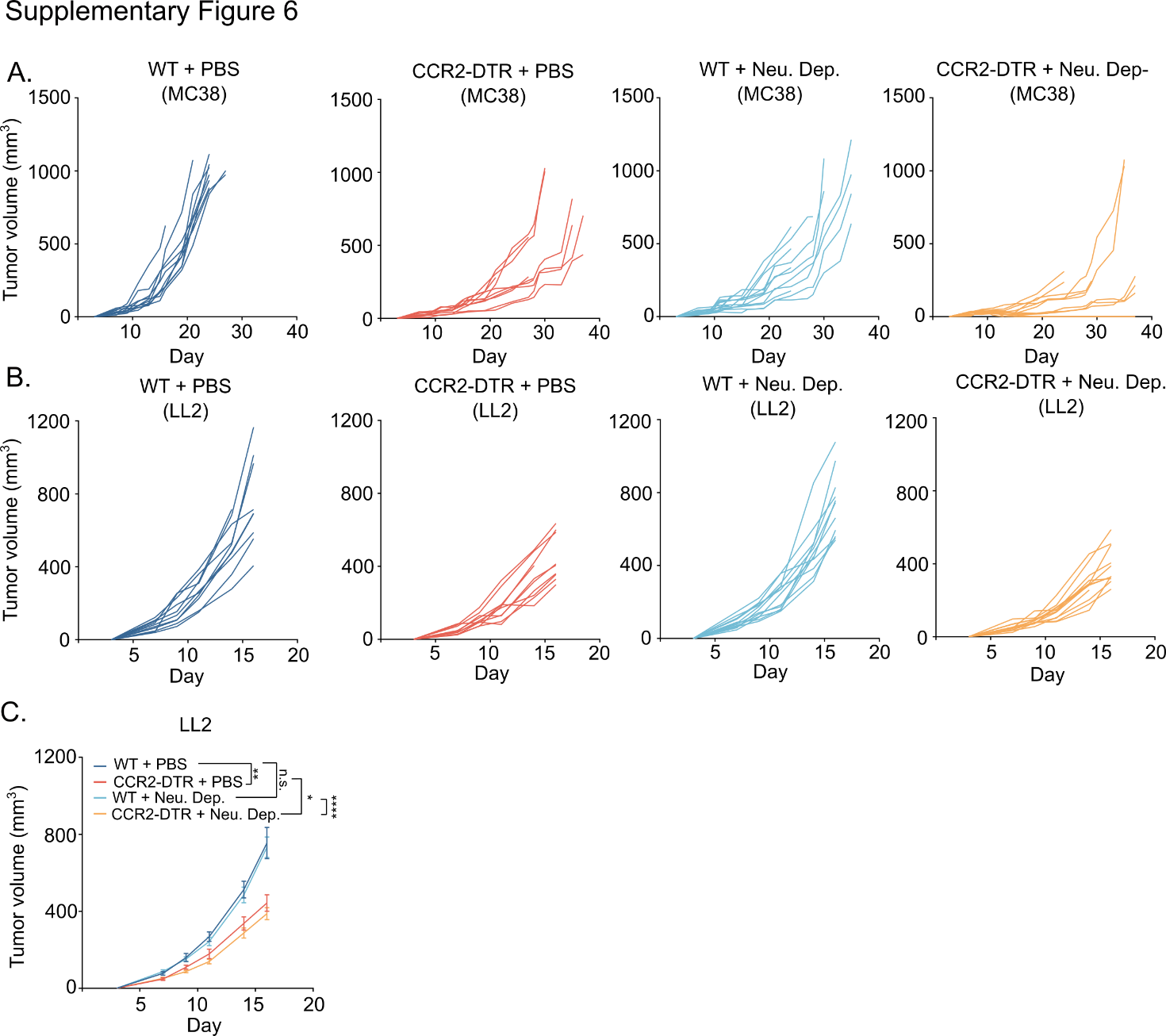

**Supplementary figure 6.**

Tumor growth kinetics of MC38 **(A)**, and LL2 **(B)** in wildtype and CCR2-DTR mice, treated with PBS or neutrophil-depleting antibodies (Neu. Dep.). Each line in tumor growth plot represents one biological replicate. **(C)** Average tumor growth of LL2 tumors in wildtype and CCR2-DTR mice, treated with PBS or neutrophil-depleting antibodies. n.s. not significant, *p<0.05, **p<0.01, ****p<0.001 according to TumGrowth software. Mean ± SEM are depicted.

Supplementary table 1. List of flow cytometry antibodies

| Vendor | Catalog number | Fluorophore | Target | Dilution factor |
| --- | --- | --- | --- | --- |
| BioLegend | 103108 | FITC | CD45 | 1:100 |
| BioLegend | 103155 | BV605 | CD45 | 1:100 |
| BioLegend | 150610 | PE | CCR2 | 1:100 |
| Miltenyi Biotech | 130-128-939 | Biotin | CCR2 | 1:50 |
| BioLegend | 128024 | AF700 | Ly6C | 1:100 |
| BioLegend | 128012 | PerCP-Cy5.5 | Ly6C | 1:100 |
| BioLegend | 127623 | APC-Cy7 | Ly6G | 1:100 |
| BioLegend | 127627 | BV241 | Ly6G | 1:100 |
| BioLegend | 127645 | BV786 | Ly6G | 1:100 |
| BioLegend | 123116 | APC | F4/80 | 1:100 |
| BioLegend | 101216 | PE-Cy7 | CD11b | 1:100 |
| BioLegend | 101226 | APC-Cy7 | CD11b | 1:100 |
| BioLegend | 123212 | APC | PDL1 | 1:100 |
| BD Biosciences | 740622 | BV650 | CD64 | 1:100 |
| BioLegend | 117306 | FITC | CD11c | 1:100 |
| Miltenyi Biotech | 130-102-892 | PE | CCR3 | 1:50 |
| BD Horizon | 562477 | PE-CF594 | CD24 | 1:100 |
| BioLegend | 100204 | FITC | CD3 | 1:100 |
| BioLegend | 116016 | PE-Cy7 | CD4 | 1:100 |
| BioLegend | 100438 | BV421 | CD4 | 1:100 |
| BioLegend | 100414 | APC-Cy7 | CD4 | 1:100 |
| BioLegend | 100712 | APC | CD8 | 1:100 |
| BD Biosciences | 566409 | BB700 | CD8 | 1:100 |
| BD Pharmigen | 551892 | PE | PD1 | 1:100 |
| BD Horizon | 562864 | PE-CF594 | NK1.1 | 1:100 |
| BioLegend | 106603 | Biotin | FasL | 1:100 |
| BioLegend | 405208 | APC-Cy7 | Streptavidin | 1:100 |
| BioLegend | 405226 | BV421 | Streptavidin | 1:100 |
| BioLegend | 165803 | PE | ARG1 | 1:100 |
| Abcam | ab90812 | FITC | MPO | 1:50 |
